# Molecular mechanisms driving divergent development of the human frontal and visual cortex during prenatal development

**DOI:** 10.1101/2024.05.15.594422

**Authors:** Gabriela Epihova, Dimitar Z. Epihov, Danyal Akarca, Duncan E. Astle

## Abstract

Key principles of structural brain organization are established very early in fetal development. The frontal cortex is an important hub for integration and control of information, and its integrity and connectivity within the wider neural system are linked to individual differences across multiple cognitive domains and neurodevelopmental conditions. Here we leveraged fetal brain transcriptomics to investigate molecular mechanisms during prenatal development that drive early differences between the two regions at the opposite poles of the physical and representational gradient of the brain - the frontal and visual cortex. We show that the frontal cortex exhibits significantly higher cumulative gene expression for pathways involved in the continued growth and maintenance of larger neurons. These pathways include the gene ontology terms of neuron development and neuronal cell body as well as glucose metabolism important in trophically supporting larger cell sizes. Whole pathways for axonal growth (axonal growth cone, microtubules, filopodia, lamellipodia) and single genes involved in circuit connectivity exhibited increased expression in the frontal cortex. In contrast, in line with the established earlier completion of neurogenesis and lower number of neurons in the anterior cortex, expression of genes involved in DNA replication was significantly lower relative to the visual cortex. We further demonstrate differential cellular composition with higher expression of marker genes for inhibitory neurons in the prenatal frontal cortex. Together, these results suggest that the cellular architecture and composition facilitates earlier connectivity in the frontal cortex which may determine its role as an integrative hub in the global brain organization.

## Introduction

Over the time-course of development, brain organization gradually develops along a unimodal-to-transmodal hierarchy with progression from domain-specific sensory processing in unimodal cortex to integrative domain-general representations from multiple sensory modalities in transmodal cortex (1,2). The frontal cortex represents the apex of this unimodal-transmodal gradient – the most anterior part of cortex which reflects the largest extent of transmodal activity (2,3). The degree of integration of the frontal cortex within the wider neural system has been implicated across higher-order cognitive domains (executive function, inhibition, working memory, planning, and reasoning) and is involved in multiple, if not all, neurodevelopmental and neuropsychiatric conditions (4-6). Consistent with this, previous studies investigating brain spatiotemporal gene expression patterns have reported that the human prefrontal cortex, and the mid-fetal developmental period, are vulnerable points for neurodevelopmental disorders (5-9).

Transcriptomic analyses of the human brain across the lifespan have revealed highly dynamic gene expression during prenatal and early postnatal development, versus comparative stability over several decades of adulthood (8, 10). However, little is known about the developmental mechanisms guiding the early emergence of frontal cortex development and the extent to which the mechanisms differ relative to other cortical areas (see 11 for an overview of frontal cortex development). There is a paucity of studies of the fetal brain organization given the technical challenges of neuroimaging *in utero* (12). However, a few studies have reported that whole brain organization already resembles that of the adult at 20–30 post-gestational weeks (13, 14), suggesting that core organisational principles of the connectome are already established in prenatal development (13, 15). Interestingly, diffusion tensor imaging (DTI) at 20 gestational weeks has shown that the fronto-medial cortex harbours increased strength of connectivity relative to the rest of the cortex (14).

One explanation of this relatively dense anterior connectivity *in utero* may be the distinct microscale properties influencing the macroscale connectome emergence. The cerebral cortex is an exceptionally complex structure, exhibiting marked regional anatomical heterogeneity (16). Specifically, the cortex harbours gradients in cytoarchitecture and timing of neurogenesis such that neurogenesis is completed earlier anteriorly compared to the posterior cortex (17-19). The shorter duration of neurogenesis in the anterior cortex is associated with decreased neuron number and density (18, 20-22), leading to anterior neurons with larger cell bodies and longer processes (23, 24). Crucially, these cytoarchitectural properties have been associated with strength of connectivity. In the macaque, brain regions with lower neural density and a larger cell size bear a higher degree of total connections (25, 26). Similarly, in the round worm *Caenorhabditis elegans* neurons generated earlier during development tend to accumulate more connections and become hubs in the adult network through the rich-get-richer principle whereby connections are preferentially forming to existing areas that already have multiple connections (27-29).

Understanding the mechanisms guiding the differential development of connectivity across the fetal cortex has widespread implications for emergence of connectivity and later manifestation of cognitive and behavioural characteristics that vary naturally across the population and are implicated across multiple neurodevelopmental conditions. Here we leveraged the BrainSpan Developing Brain atlas – a gene expression brain atlas covering the prenatal period – to investigate the transcriptomic signatures of cytoarchitectural and connectivity differences along the anterior-posterior cortical gradient during early-to-mid prenatal development. Specifically, we focused on the differential development of the two regions situated at the opposite poles of the cortical gradient - the frontal and visual cortex. Finally, to further characterise early functional differences between the frontal and visual cortex we asked whether the composition of cell types differs by comparing expression levels of cell type markers in the bulk transcriptome data.

## Results

### Enhanced axon growth cone processes in the frontal compared to visual cortex

We first compared the cumulative expression of genes for each gene ontology (GO) term between frontal and visual cortical regions in samples from 12 to 24 pgws (see Materials and Methods). GO terms that were differentially expressed can be found in Supplementary File 1 and are available at https://osf.io/s4qb2/. The visual cortex was enriched for the processes of DNA replication (p<0.001) indicating a higher neuron cycle rate resulting in more neurons (**Fig 1a, in blue**). The visual cortex also showed an increased expression of genes for negative regulation of neurogenesis (p=0.003), suggesting that in the visual cortex there is a delay in the cell cycle exit, thus prolonging neurogenesis, consistent with the previously reported anterior-posterior cortical gradient of neurogenesis timing (18-19). In contrast, the frontal cortex was enriched for the pathways of neuron development (p=0.033), neuronal cell body (p=0.008), cytoplasm (p=0.010), mitochondrion (p=0.013), cytoskeleton (p<0.001), and glucose metabolic process (p=0.007), (**Fig 1a, in red**). We interpret these findings as hallmarks for larger-celled neuronal population anteriorly, that are energetically supported by mirroring increases in central carbohydrate metabolism. In accordance with these results, cumulative expression of genes implicated in ATP synthase complex (p=0.007) and oxidative phosphorylation (p=0.003) were also increased in the frontal cortex. These results are compatible with previous evidence in the adult primate cortex of larger cell bodies and longer processes in the anterior vs posterior cortical poles (30). Further, our findings are consistent with the increased metabolic activity seen in the frontal cortex relative to the rest of the human brain in adulthood (31).

**Figure 1.**
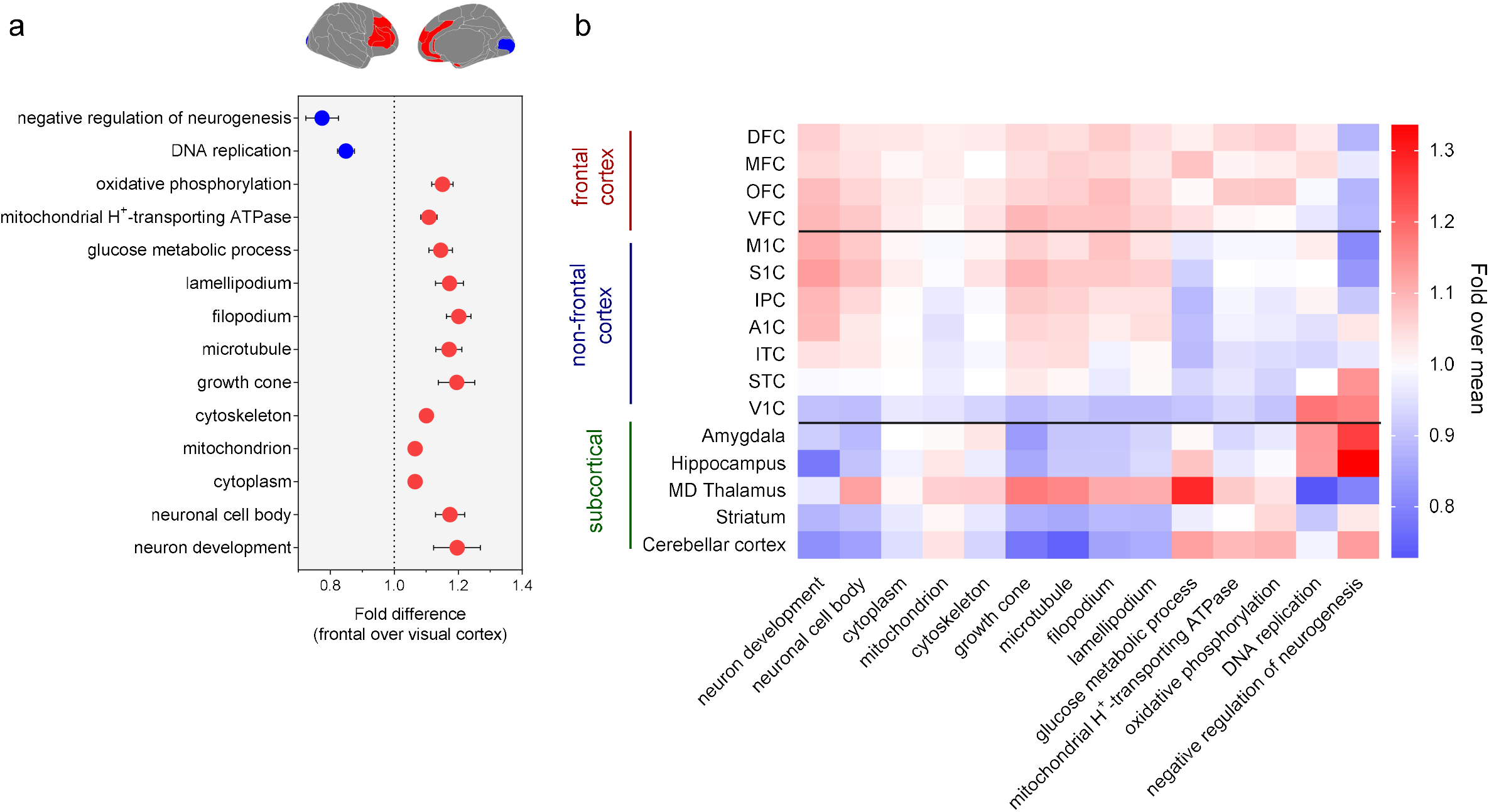
**a**. Fold difference (frontal/visual cortex ratio in expression) in cumulative expression of selected pathways between the frontal and visual cortex. **b**. Expression (fold over mean) of selected pathways across all cortical and subcortical brain regions. Fold over mean expression is the expression value of each region divided by the average expression across all regions for each pathway.

During brain development emerging connectivity occurs through communication between guidance cues and the cell’s growing tip of the axon – the growth cone. Growth cones at the tip of advancing axons pull themselves and their axons forward. Structurally the growth cone consists of microtubules, F-actin network and bundles, filopodia, and lamellipodia (32-34).

Filopodia and lamellipodia are dynamic structures - they can form, extend, or withdraw within seconds to minutes in response to environmental cues such as chemoattractants and chemorepellents. This is achieved through rearrangements of microtubule and actin networks of the growth cone (35-37). Microtubules are fibres composed of α-tubulin and β-tubulin with the majority of microtubules in a neuronal cell residing along the axon and dendrites (38, 39). Critically, we show that these processes (growth cone (p=0.014), microtubule (p=0.004), filopodium (p<0.001) and lamellipodium (p=0.006) that are directly related to emergent connectivity had increased cumulative expression in regions within the frontal cortex relative to the visual cortex. This further suggests that cortical cytoarchitecture appears to be associated with the organization of early connections.

To examine whether the observed effects are limited to the frontal-visual contrast, we examined gene expression of these selected pathways across all available cortical and subcortical regions. The expression of these pathways showed an anterior-posterior spatial gradient, with highest expression in fronto-parietal regions, followed by temporal cortex, and lowest expression in posterior visual cortex (**Fig 1b**). It is of note that the expression of pathways relating to cell development and size (neuron development, neuronal cell size, cytoplasm, and cytoskeleton) was similar in parietal cortex (motor and somatosensory regions) to that of the frontal cortex. It has been previously shown that although neural density follows a general anterior-posterior gradient, a region proximal to the motor cortex is an outlier in the sense that it harbours sparser neural density even relative to the frontal cortex (20). One additional observation is that the enriched pathways in the frontal relative to visual cortex generally had very low expression in the rest of the subcortical structures but had the highest expression in the medio-dorsal (MD) thalamus (**Fig 1b**). In particular, consistent with its role of a relay station, the MD thalamus was over-expressing genes involved in the establishment of axonal connectivity (growth cone, microtubules, filopodia, lamellipodia) and had the highest glucose metabolism relative to the rest of brain regions.

### Higher expressions of genes involved in axonal growth in the frontal cortex

Next, we carried out multiple anterior vs. posterior contrasts to assess expression differences on a single gene-level. Genes that were differentially expressed can be found in Supplementary File 2 and are available at https://osf.io/s4qb2/. Our analysis confirmed previous reports of frontally-enriched genes (40). In line with our research goal, we focused on regional variation in genes previously implicated in axonal development, growth, guidance, and circuit connectivity. There was a significantly higher expression in the frontal compared to visual cortex for *GPM6A* (p=0.001) which is involved in filopodia formation and motility, promoting neurite outgrowth and axon elongation during development (41, 42). Relatedly, expression of Fascin Actin-Bundling Protein 1 (*FSCN1)* was enriched anteriorly (p=0.008). Fascin-dependent actin bundling is essential for axonal growth by organizing F-actin into parallel bundles thus promoting the formation of actin-based cellular protrusions such as filopodia in the growth cone (43). Frontally, there were significantly higher levels of brain derived neural factor (*BDNF*), (p=0.008) which promotes growth and guidance of axons. Axon growth cones exposed to *BDNF* in culture respond to extracellular gradients of *BDNF* (44). Expression levels were also higher for *TUBA1A* (p=0.002), *TUBB2B* (p=0.017), *TUBB3* (p=0.005), which provide instructions for making version of α-tubulin, and β-tubulin that form microtubules. Further, there was higher expression of *NTN1* (p<0.0001), *ROBO1* (p<0.0001), *NRP2* (p<0.0001), *GAP43* (p=0.012), *SCGN* (p<0.001) and *SNAP25* (p=0.003) which contribute to axon growth and guidance (45-50). Overall, these axonal growth genes had higher expression in the frontal cortex not only relative to V1, but to other more posterior cortical regions (**Fig. 2**).

**Figure 2.**
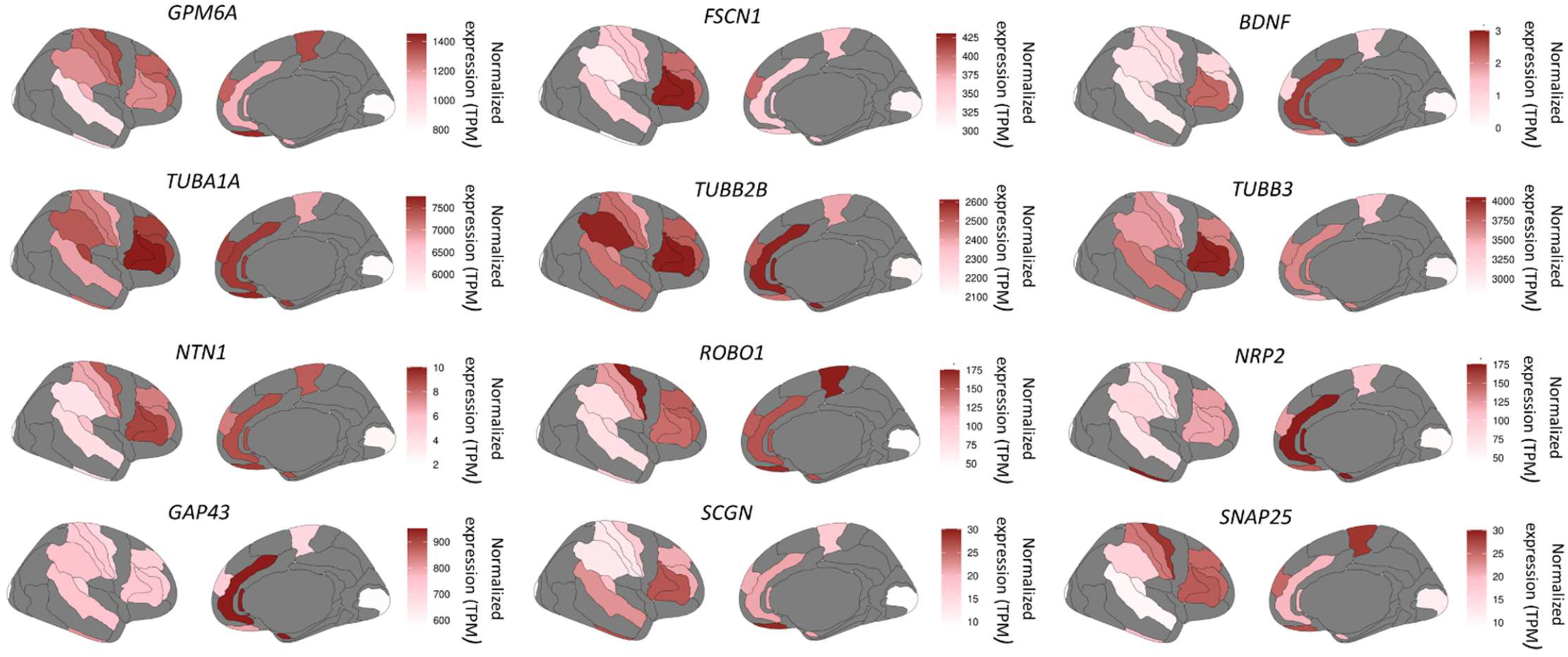
Anterior-posterior gradient in normalized expression levels (TPM) of genes linked to axonal development, growth, guidance, and circuit connectivity during early-to-mid prenatal development (12-24 pgwk). Gene expression is mapped onto the adult Brodmann atlas for visualization.

### Inhibitory neurons are enriched in the frontal relative to the visual cortex

Next, we investigated how the proportion of cell types differs between the frontal and visual cortex by contrasting the expression levels of marker genes for the different cell types (see Materials and Methods). Marker genes are a set of genes with highly enriched expression in a particular cell type (**Fig 3a**), and relative differences in their expression can be used to estimate differences in cell types across bulk RNAseq samples (51-53). Specifically, we explored expression differences in marker genes in the bulk RNA samples from the BrainSpan Developing Brain atlas for early fetal, late fetal, and postnatal excitatory neurons, fetal inhibitory neurons, medial ganglionic eminence-derived inhibitory neurons (IN-MGE), caudal ganglionic eminence-derived inhibitory neurons (IN-CGE), oligodendrocyte progenitor cells (OPC), oligodendrocytes, astrocytes, microglia, radial glia, intermittent progenitor cells (IPC), endothelial cells, pericytes, and vascular smooth muscle cells (VSMC).

**Figure 3.**
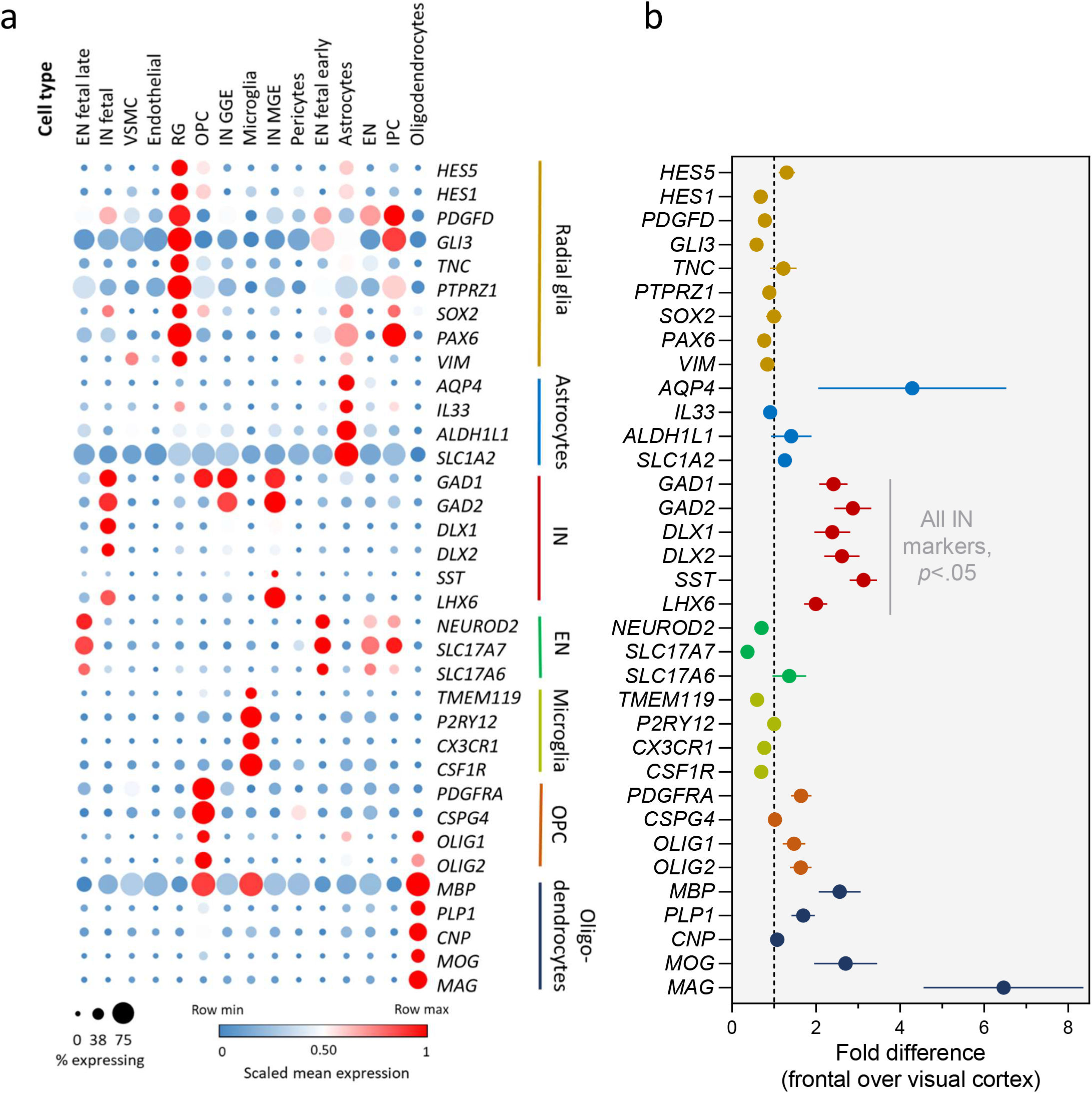
**a**. Marker genes show distinct expression for specific cell types; **b**. Higher expression of marker genes for inhibitory neurons (*GAD1, GAD2, DLX1, DLX2, SST, LHX6*) in frontal cortex relative to visual cortex during early-to-mid prenatal development (12-24pgwk).

We observed higher expression of inhibitory neurons in prenatal frontal cortex relative to the visual cortex (*GAD1* p=0.003, *GAD2* p=0.003, *DLX1* p=0.019, *DLX2* p=0.010, *SST* p<0.001, *LHX6* p=0.005), (**Fig 3b**). There were no consistent significant differences across other cell type maker genes at p<0.05.

The higher expression in the frontal cortex of marker genes for inhibitory neurons was confined to the prenatal period when inhibitory neurons have an excitatory function (54-57), with little differences in post-natal expression (except *SST*) (**Fig. 4**). The pattern of higher expression in frontal versus posterior visual region holds valid both for markers whose expression was highest post-natally (*GAD1, GAD2* and *SST*) as well as inhibitory markers with highest expression in the prenatal period (*DLX1, DLX2* and *LHX6)*.

**Figure 4.**
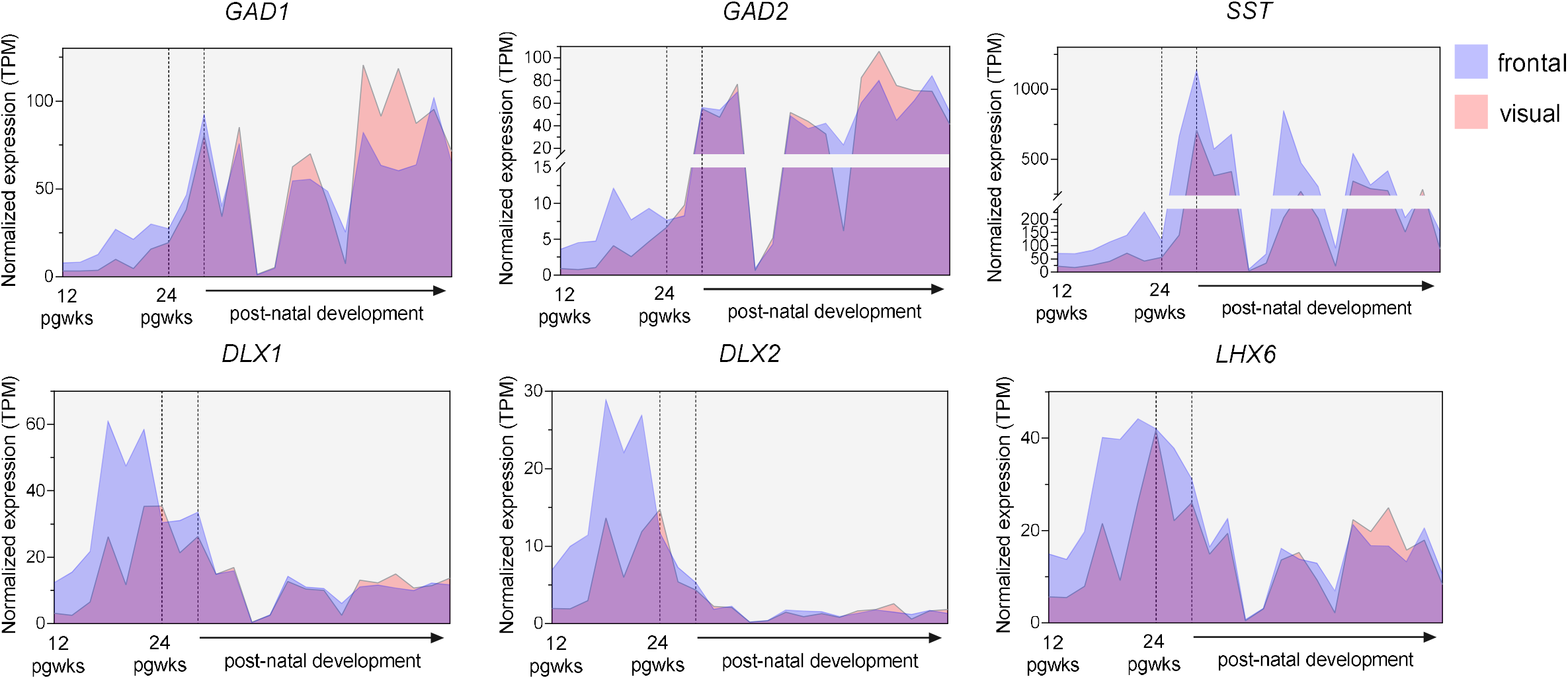
Expression levels of inhibitory neuron markers from 12 pgwks to adulthood (40 years) in the frontal and visual cortex. Inhibitory markers had higher expression levels in the frontal cortex during early-to-mid fetal development (12-24 pgwks), and this was limited to the prenatal period, with no difference in expression in postnatal development (except *SST*).

## Discussion

Prenatal development, arguably the most important period for brain development and vulnerability to neurodevelopmental disorders, is also the most challenging to study. Here we leveraged prenatal gene expression to investigate the transcriptomic signatures of early frontal cortex development. In particular, we explored early molecular mechanisms underlying cytoarchitectural and connectivity differences between two regions situated at the opposite poles of the physical space and representational continuum of the brain - the frontal and visual cortex. Overall, our results highlight the importance of early emergence of differences in microscale cytoarchitectural properties to the vastly divergent functional properties of the two regions in the adult brain – the frontal cortex acts as a dimensionality reduction, integrative hub whereas the visual cortex more faithfully maps to the sources of its activation. During prenatal development, the frontal cortex already harbours larger cellular processes supported by higher glucose metabolism. This is consistent with the previously reported anterior-to-posterior cortical gradient in the monkey cortex: sparser cell density and greater neuronal size in the frontal cortex, and more dense but smaller cellular process in the visual cortex (18, 20-22).

What are the mechanisms leading to sparser but larger neurons in the frontal cortex? One mechanism resulting in lower density of neurons in the frontal cortex is through an anterior-posterior gradient in neurogenesis duration which could generate this disparity in neuron number (30). We found decreased expression for DNA replication in the anterior brain, reflecting a lower rate of neurogenesis. One manner in which neurogenesis is controlled is through interneuron GABA signalling. We found higher expression of inhibitory neuron marker genes in the frontal cortex, which was limited to the prenatal period. GABA (ɣ-aminobutyric acid) is the main inhibitory neurotransmitter in the brain and acts primarily by binding to GABA_A_ or GABA_B_ receptors which is critical for normal development of cortical circuits (58). While GABA interneurons are inhibitory in the postnatal brain, they have an excitatory function in embryonic cortical development (54-57), with a shift towards the well-known inhibitory function at around the first postnatal week in humans (59). During prenatal development, inhibitory neurons can release GABA and activate GABA receptors on radial glia, depolarizing these progenitors and decreasing their proliferation via activation of voltage-gated Ca^2+^ channels thus negatively regulating DNA synthesis (56, 60). In addition to regulating proliferation, GABA_A_ receptor activation has been shown to promote neurite outgrowth and maturation of GABA interneurons (61). One pathway through which these neurotrophic effects might be achieved is by enhanced BDNF expression and release from target neurons (62, 63), which is in line with our results of increased BDNF expression in the frontal prenatal cortex.

Key genes and processes (growth cone, microtubules, filopodia, lamellipodia) involved in axonal growth and migration were enhanced in the frontal cortex, relative to the visual cortex. This is in line with previous evidence of the frontal cortex as an early connectivity hub *in utero* (14). The link between the cytoarchitectural properties of the frontal cortex (larger neurons size and sparser cell density) and its increased connectivity has been established in the macaque brain (25, 26). Our results demonstrate this association in the human brain by showing that the molecular pathways supporting larger cell size and axon extension are enhanced in the frontal cortex and this relationship is established very early on in prenatal development. It is of note that present mathematical models concerning axonal outgrowth patterns (64-65) do not consider these cell-specific factors across spatial axes, offering potentially new computational avenues to examine the formation of early prenatal connectivity in the human brain with greater specificity.

The finding that the frontal cortex has higher connectivity strength in the prenatal brain is somewhat counterintuitive to the fact that it is the last region in the brain to complete myelination of its connections – a process taking place well into adulthood (2). We propose that it is precisely the combination of early over-connectivity and later myelination that enables the frontal cortex to serve its function as an integrative hub supporting increasingly abstract levels of representation throughout the human lifespan. There are two main properties that an integrative hub needs to satisfy in order to perform integration of information: 1) connect to multiple regions, and 2) keep long-term plasticity of these connections to incorporate changing inputs at differing rates from other regions as they mature. To this end, our results demonstrate that the frontal cortex is enriched in molecular mechanisms for axon extension required for early connectivity. In line with the second property, the frontal cortex develops myelination in its connections last (2, 66). Although, myelination contributes to faster information transfer and therefore serves a more efficient communication, it is also one of the processes that inhibits brain plasticity and closes critical periods (67, 68). Thus, by setting up early connections and allowing for these connections to be malleable (less myelinated) over a longer time, the frontal cortex may flexibly integrate information.

Using the same dataset it has been demonstrated that across expression of all human genes, the frontal cortex (MFC, OFC, DFC and VFC) formed a distinct cluster, and together with M1C, and S1C areas formed a separate larger cluster distinguishable from another large cluster composed of IPC, STC, and A1C and ITC. The visual cortex (V1) was the most transcriptionally divergent among the analysed cortical areas (6, 10). This is consistent with the subset of pathways we currently analysed. However, we show that the majority of pathways and genes which differed significantly between frontal and visual cortex also exhibited an anterior-to-posterior gradient in expression. This suggests that the differences between the frontal and visual cortex are also present with other brain regions, although to a lesser extent. One interesting exception was the process of glucose metabolism, mitochondrion H^+^-transporting ATP synthase complex, and oxidative phosphorylation which did not show a gradient in expression, but rather selectively high expression only in the frontal cortex across all cortical regions sampled. This cannot be accounted solely on the need to support larger cell bodies, or axonal circuit formation as the difference in expression for these processes were not so prominent between the frontal cortex and the rest of the cortex. Other processes requiring higher glucose metabolism in the prenatal frontal cortex should be explored in future investigations.

In conclusion, we showed that the frontal cortex exhibits significantly higher cumulative gene expression for pathways involved in the continued growth and maintenance of fewer but larger neurons. The lower number of neurons in the frontal cortex might result through an earlier completion of neurogenesis and GABA-induced inhibition of neural progenitors proliferation. Additionally, pathways for axonal growth and single genes involved in circuit connectivity exhibited increased expression in the frontal cortex. Together, these results suggest that early cell architecture and composition facilitates increased connectivity in the frontal cortex which determines its role as an integrative hub in the global brain organization.

## Materials and Methods

### Transcriptomics data

RNAseq data was obtained from the publicly available BrainSpan Developing Brain atlas (https://www.brainspan.org). The data used in the current analyses covered gene expression from 11 cortical: dorsal frontal cortex (DFC), medial frontal cortex (MFC), orbito-frontal cortex (OFC), ventral frontal cortex (VFC), motor cortex (M1C), somato-sensory cortex (S1C), auditory cortex (A1C), inferior parietal cortex (IPC), superior temporal cortex (STC), inferior temporal cortex (ITC), visual cortex (V1C) and 5 sub-cortical regions: amygdala, cerebellum, hippocampus, striatum, dorsal thalamus. Data from corresponding regions in the left and right hemispheres were pooled together. The obtained gene expression data were in reads per kilobase per million (RPKM) values. To allow normalized comparisons across samples, RPKM values were converted to transcripts per million (TPM) according to the formula:

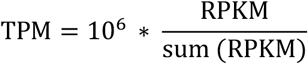

Genes were further annotated to functional pathways by linking their gene IDs to pre-populated list of associated gene ontology (GO) terms available on the PANTHER GO server (69). To obtain information on the total number of transcripts involved in a particular pathway (as defined by GO ontology), the relative TPM expression of each of the genes within a pathway were added together as specified below:

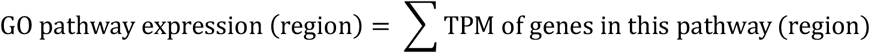

Compared to typical GO enrichment analyses which are independent on exact gene expression levels, our derived pathway expression allows us to explore the additive effects of all genes contributing to a certain pathway and test for differences in cumulative expression between regions.

We validated our approach by showing that the cumulative gene expression for selected GO pathways follows well-established patterns in brain development across the life span (**Supp. Fig 1**).

### Early-to-mid prenatal anterior – posterior regions contrast

A Uniform Manifold Approximation and Projection (UMAP) on the gene expression data was performed for dimensionality reduction of the 18,880 GO terms (**Supp. Fig 2**). The UMAP results indicated that at 37 weeks the expression patterns transition to a distinct state compared to earlier fetal and *ex utero* expression, replicating the previously reported transcriptomic transition beginning during late fetal development (8, 10). Because of this we excluded data from 37 pgwk and constrained our analyses on the remaining early-to-mid prenatal samples from 12 to 24 post-gestational weeks (pgwk). This resulted in 13 donor samples (6 female, 7 male): 3 donors at 12 pgwk, 3 donors at 13 pgwk, 3 donors at 16 pgwk, 1 donor at 17 pgwk, 1 donor at 19 pgwk, 1 donor at 21 pgwk, 1 donor at 24 pgwk. We focused the comparison between the most anterior and posterior regions of the cortex – the frontal and visual cortex. Gene expression from the frontal cortex included expression from the medial, dorsal, ventral, and occipital frontal regions (MFC, DFC, VFC and OFC - Broadman areas 9, 11, 32, 33, 34, 44, 45, 46), which was contrasted with gene expression from the posterior brain (V1 cortex - Broadman area 17). Donors without expression data from V1 cortex and at least one frontal region were removed from the comparison to ensure no individual differences bias in expression.

### Early-to-mid prenatal anterior – posterior regions cell type enrichment analysis

First, we cross-referenced known brain cell type marker genes with a developmentally-relevant single-cell RNAseq data which were derived from frontal cortex tissue across developmental timepoints from early fetal to adulthood (70). Expression of maker genes were referenced across a set of pre-defined neuronal cell types: early fetal excitatory neurons (EN fetal early), late fetal excitatory neurons (EN fetal late), post-natal excitatory neurons (EN), fetal inhibitory neurons (IN fetal), medial ganglionic eminence-derived inhibitory neurons (IN-MGE), caudal ganglionic eminence-derived inhibitory neurons (IN-CGE), oligodendrocyte progenitor cells (OPC), oligodendrocytes, astrocytes, microglia, radial glia, intermittent progenitor cells (IPC), endothelial cells, pericytes, and vascular smooth muscle cells (VSMC). After ensuring that the marker genes are expressed uniquely in a cell-type, we investigated whether expression of these markers in the bulk RNA samples from the BrainSpan Developing Brain atlas differed between the frontal and visual regions.

## Acknowledgements

G.E, D.A and D.E.A were supported by an Opportunity Award from the James S. McDonnell Foundation. D.E.A was supported by the Medical Research Council program grant MC-A0606-5PQ41. D.E.A and D.A were also supported by the Templeton World Charity Foundation, Inc. (funder DOI 501100011730) under grant TWCF-2022-30510. All research at the Department of Psychiatry in the University of Cambridge is supported by the National Institute for Health and Care Research Cambridge Biomedical Research Centre (NIHR203312) and the NIHR Applied Research Collaboration East of England. We also thank the authors of the publicly available databases used in this study, without whom this work would not have been possible.

## Data availability

Code and results from the current analyses are publicly available at https://osf.io/s4qb2/

## References

1. Huntenburg, J. M., Bazin, P. L., & Margulies, D. S. (2018). Large-scale gradients in human cortical organization. Trends in cognitive sciences, 22(1), 21–31.

2. Sydnor, V. J., Larsen, B., Seidlitz, J., Adebimpe, A., Alexander-Bloch, A. F., Bassett, D. S., … & Satterthwaite, T. D. (2023). Intrinsic activity development unfolds along a sensorimotor–association cortical axis in youth. Nature Neuroscience, 26(4), 638–649.

3. Sydnor, V. J., Larsen, B., Bassett, D. S., Alexander-Bloch, A., Fair, D. A., Liston, C., … & Satterthwaite, T. D. (2021). Neurodevelopment of the association cortices: Patterns, mechanisms, and implications for psychopathology. Neuron, 109(18), 2820–2846.

4. Schubert, D., Martens, G. J. M., & Kolk, S. M. (2015). Molecular underpinnings of prefrontal cortex development in rodents provide insights into the etiology of neurodevelopmental disorders. Molecular psychiatry, 20(7), 795–809.

5. Gulsuner, S., Walsh, T., Watts, A. C., Lee, M. K., Thornton, A. M., Casadei, S., … & McClellan, J. M. (2013). Spatial and temporal mapping of de novo mutations in schizophrenia to a fetal prefrontal cortical network. Cell, 154(3), 518–529.

6. Willsey, A. J., Sanders, S. J., Li, M., Dong, S., Tebbenkamp, A. T., Muhle, R. A., … & Sestan, N. (2013). Coexpression networks implicate human midfetal deep cortical projection neurons in the pathogenesis of autism. Cell, 155(5), 997–1007.

7. Werling, D. M., Pochareddy, S., Choi, J., An, J. Y., Sheppard, B., Peng, M., … & Sestan, N. (2020). Whole-genome and RNA sequencing reveal variation and transcriptomic coordination in the developing human prefrontal cortex. Cell reports, 31(1).

8. Li, M., Santpere, G., Imamura Kawasawa, Y., Evgrafov, O. V., Gulden, F. O., Pochareddy, S., … & Sestan, N. (2018). Integrative functional genomic analysis of human brain development and neuropsychiatric risks. Science, 362(6420), eaat7615.

9. Chang, J., Gilman, S. R., Chiang, A. H., Sanders, S. J., & Vitkup, D. (2015). Genotype to phenotype relationships in autism spectrum disorders. Nature neuroscience, 18(2), 191–198.

10. Kang, H. J., Kawasawa, Y. I., Cheng, F., Zhu, Y., Xu, X., Li, M., … & Šestan, N. (2011). Spatio-temporal transcriptome of the human brain. Nature, 478(7370), 483–489.

11. Kolk, S. M., & Rakic, P. (2022). Development of prefrontal cortex. Neuropsychopharmacology, 47(1), 41–57.

12. Christiaens, D., Slator, P. J., Cordero-Grande, L., Price, A. N., Deprez, M., Alexander, D. C., … & Hutter, J. (2019). In utero diffusion MRI: challenges, advances, and applications. Topics in Magnetic Resonance Imaging, 28(5), 255–264.

13. Ball, G., Aljabar, P., Zebari, S., Tusor, N., Arichi, T., Merchant, N., … & Counsell, S. J. (2014). Rich-club organization of the newborn human brain. Proceedings of the National Academy of Sciences, 111(20), 7456–7461.

14. Song, L., Mishra, V., Ouyang, M., Peng, Q., Slinger, M., Liu, S., & Huang, H. (2017). Human fetal brain connectome: structural network development from middle fetal stage to birth. Frontiers in neuroscience, 11, 561.

15. Yap, P. T., Fan, Y., Chen, Y., Gilmore, J. H., Lin, W., & Shen, D. (2011). Development trends of white matter connectivity in the first years of life. PloS one, 6(9), e24678.

16. Rakic, P. (2008). Confusing cortical columns. Proceedings of the National Academy of Sciences, 105(34), 12099–12100.

17. Rakic, P. (1974). Neurons in rhesus monkey visual cortex: systematic relation between time of origin and eventual disposition. Science, 183(4123), 425–427.

18. Charvet, C. J., & Finlay, B. L. (2014). Evo-devo and the primate isocortex: the central organizing role of intrinsic gradients of neurogenesis. Brain Behavior and Evolution, 84(2), 81–92.

19. Charvet, C. J., Cahalane, D. J., & Finlay, B. L. (2015). Systematic, cross-cortex variation in neuron numbers in rodents and primates. Cerebral Cortex, 25(1), 147–160.

20. Collins, C. E., Turner, E. C., Sawyer, E. K., Reed, J. L., Young, N. A., Flaherty, D. K., & Kaas, J. H. (2016). Cortical cell and neuron density estimates in one chimpanzee hemisphere. Proceedings of the National Academy of Sciences, 113(3), 740–745.

21. Collins, C. E., Airey, D. C., Young, N. A., Leitch, D. B., & Kaas, J. H. (2010). Neuron densities vary across and within cortical areas in primates. Proceedings of the National Academy of Sciences, 107(36), 15927–15932.

22. Cahalane, D. J., Charvet, C. J., & Finlay, B. L. (2012). Systematic, balancing gradients in neuron density and number across the primate isocortex. Frontiers in neuroanatomy, 6.

23. Beul, S. F., Barbas, H., & Hilgetag, C. C. (2017). A predictive structural model of the primate connectome. Scientific reports, 7(1), 43176.

24. Beul, S. F., & Hilgetag, C. C. (2019). Neuron density fundamentally relates to architecture and connectivity of the primate cerebral cortex. NeuroImage, 189, 777–792.

25. Scholtens, L. H., Schmidt, R., de Reus, M. A., & van den Heuvel, M. P. (2014). Linking macroscale graph analytical organization to microscale neuroarchitectonics in the macaque connectome. Journal of Neuroscience, 34(36), 12192–12205.

26. van den Heuvel, M. P., Scholtens, L. H., Barrett, L. F., Hilgetag, C. C., & de Reus, M. A. (2015). Bridging cytoarchitectonics and connectomics in human cerebral cortex. Journal of Neuroscience, 35(41), 13943–13948.

27. Varier, S., & Kaiser, M. (2011). Neural development features: spatio-temporal development of the Caenorhabditis elegans neuronal network. PLoS computational biology, 7(1), e1001044.

28. Towlson, E. K., Vértes, P. E., Ahnert, S. E., Schafer, W. R., & Bullmore, E. T. (2013). The rich club of the C. elegans neuronal connectome. Journal of Neuroscience, 33(15), 6380–6387.

29. Oldham, S., Ball, G., & Fornito, A. (2022). Early and late development of hub connectivity in the human brain. Current opinion in psychology, 44, 321–329.

30. Finlay, B. L., & Uchiyama, R. (2015). Developmental mechanisms channeling cortical evolution. Trends in neurosciences, 38(2), 69–76.

31. Castrillon, G., Epp, S., Bose, A., Fraticelli, L., Hechler, A., Belenya, R., … & Riedl, V. (2023). An energy costly architecture of neuromodulators for human brain evolution and cognition. Science advances, 9(50), eadi7632.

32. Letourneau, P. C. (1983). Axonal growth and guidance. Trends in Neurosciences, 6, 451–455.

33. Sanes, D. H., Reh, T. A., & Harris, W. A. (2011). Development of the nervous system. Academic press.

34. Huber, A. B., Kolodkin, A. L., Ginty, D. D., & Cloutier, J. F. (2003). Signaling at the growth cone: ligand-receptor complexes and the control of axon growth and guidance. Annual review of neuroscience, 26(1), 509–563.

35. Tanaka, E., & Sabry, J. (1995). Making the connection: cytoskeletal rearrangements during growth cone guidance. Cell, 83(2), 171–176.

36. Desai, A., & Mitchison, T. J. (1997). Microtubule polymerization dynamics. Annual review of cell and developmental biology, 13(1), 83–117.

37. Dent, E. W., Callaway, J. L., Szebenyi, G., Baas, P. W., & Kalil, K. (1999). Reorganization and movement of microtubules in axonal growth cones and developing interstitial branches. Journal of Neuroscience, 19(20), 8894–8908.

38. Janke, C., & Magiera, M. M. (2020). The tubulin code and its role in controlling microtubule properties and functions. Nature Reviews Molecular Cell Biology, 21(6), 307–326.

39. Baas, P. W., Rao, A. N., Matamoros, A. J., & Leo, L. (2016). Stability properties of neuronal microtubules. Cytoskeleton, 73(9), 442–460.

40. Johnson, M. B., Kawasawa, Y. I., Mason, C. E., Krsnik, Ž., Coppola, G., Bogdanović, D., … & Šestan, N. (2009). Functional and evolutionary insights into human brain development through global transcriptome analysis. Neuron, 62(4), 494–509.

41. León, A., Aparicio, G. I., & Scorticati, C. (2021). Neuronal glycoprotein M6a: an emerging molecule in chemical synapse formation and dysfunction. Frontiers in Synaptic Neuroscience, 13, 661681.

42. Honda, A., Ito, Y., Takahashi-Niki, K., Matsushita, N., Nozumi, M., Tabata, H., … & Igarashi, M. (2017). Extracellular signals induce glycoprotein M6a clustering of lipid rafts and associated signaling molecules. Journal of Neuroscience, 37(15), 4046–4064.

43. Nozumi, M., Nakatsu, F., Katoh, K., & Igarashi, M. (2017). Coordinated movement of vesicles and actin bundles during nerve growth revealed by super resolution microscopy. Cell Rep 18: 2203–2216.

44. Cohen-Cory, S., Kidane, A. H., Shirkey, N. J., & Marshak, S. (2010). Brain-derived neurotrophic factor and the development of structural neuronal connectivity. Developmental neurobiology, 70(5), 271–288.

45. Kennedy, T. E., Serafini, T., de la Torre, J., & Tessier-Lavigne, M. (1994). Netrins are diffusible chemotropic factors for commissural axons in the embryonic spinal cord. Cell, 78(3), 425–435.

46. Kidd, T., Brose, K., Mitchell, K. J., Fetter, R. D., Tessier-Lavigne, M., Goodman, C. S., & Tear, G. (1998). Roundabout controls axon crossing of the CNS midline and defines a novel subfamily of evolutionarily conserved guidance receptors. Cell, 92(2), 205–215.

47. Giger, R. J., Cloutier, J. F., Sahay, A., Prinjha, R. K., Levengood, D. V., Moore, S. E., … & Geppert, M. (2000). Neuropilin-2 is required in vivo for selective axon guidance responses to secreted semaphorins. Neuron, 25(1), 29–41.

48. Skene, J. P., Jacobson, R. D., Snipes, G. J., McGuire, C. B., Norden, J. J., & Freeman, J. A. (1986). A protein induced during nerve growth (GAP-43) is a major component of growth-cone membranes. Science, 233(4765), 783–786.

49. Liu, Z., Tan, S., Zhou, L., Chen, L., Liu, M., Wang, W., … & Jia, D. (2023). SCGN deficiency is a risk factor for autism spectrum disorder. Signal Transduction and Targeted Therapy, 8(1), 3.

50. Osen-Sand, A., Catsicas, M., Staple, J. K., Jones, K. A., Ayala, G., Knowles, J., … & Catsicas, S. (1993). Inhibition of axonal growth by SNAP-25 antisense oligonucleotides in vitro and in vivo. Nature, 364(6436), 445–448.

51. Jew, B., Alvarez, M., Rahmani, E., Miao, Z., Ko, A., Garske, K. M., … & Halperin, E. (2020). Accurate estimation of cell composition in bulk expression through robust integration of single-cell information. Nature communications, 11(1), 1971.

52. Seidlitz, J., Nadig, A., Liu, S., Bethlehem, R. A., Vértes, P. E., Morgan, S. E., … & Raznahan, A. (2020). Transcriptomic and cellular decoding of regional brain vulnerability to neurogenetic disorders. Nature communications, 11(1), 3358.

53. Mandal, A. S., Romero-Garcia, R., Hart, M. G., & Suckling, J. (2020). Genetic, cellular, and connectomic characterization of the brain regions commonly plagued by glioma. Brain, 143(11), 3294–3307.

54. Murata, Y., & Colonnese, M. T. (2020). GABAergic interneurons excite neonatal hippocampus in vivo. Science advances, 6(24), eaba1430.

55. Ben-Ari, Y. (2002). Excitatory actions of gaba during development: the nature of the nurture. Nature Reviews Neuroscience, 3(9), 728–739.

56. Wang, D. D., & Kriegstein, A. R. (2009). Defining the role of GABA in cortical development. The Journal of physiology, 587(9), 1873–1879.

57. Owens, D. F., Boyce, L. H., Davis, M. B., & Kriegstein, A. R. (1996). Excitatory GABA responses in embryonic and neonatal cortical slices demonstrated by gramicidin perforated-patch recordings and calcium imaging. Journal of Neuroscience, 16(20), 6414–6423.

58. Peerboom, C., & Wierenga, C. J. (2021). The postnatal GABA shift: a developmental perspective. Neuroscience & Biobehavioral Reviews, 124, 179–192.

59. Kilb, W. (2012). Development of the GABAergic system from birth to adolescence. The Neuroscientist, 18(6), 613–630.

60. LoTurco, J. J., Owens, D. F., Heath, M. J., Davis, M. B., & Kriegstein, A. R. (1995). GABA and glutamate depolarize cortical progenitor cells and inhibit DNA synthesis. Neuron, 15(6), 1287–1298.

61. Owens, D. F., & Kriegstein, A. R. (2002). Is there more to GABA than synaptic inhibition?. Nature Reviews Neuroscience, 3(9), 715–727.

62. Marty, S., Berninger, B., Carroll, P., & Thoenen, H. (1996). GABAergic stimulation regulates the phenotype of hippocampal interneurons through the regulation of brain-derived neurotrophic factor. Neuron, 16(3), 565–570.

63. Berninger, B., Marty, S., Zafra, F., Berzaghi, M. D. P., Thoenen, H., & Lindholm, D. (1995). GABAergic stimulation switches from enhancing to repressing BDNF expression in rat hippocampal neurons during maturation in vitro. Development, 121(8), 2327–2335.

64. Song, H. F., Kennedy, H., & Wang, X. J. (2014). Spatial embedding of structural similarity in the cerebral cortex. Proceedings of the National Academy of Sciences, 111(46), 16580–16585.

65. Liu, Y., Seguin, C., Betzel, R. F., Akarca, D., Di Biase, M. A., & Zalesky, A. (2024). A generative model of the connectome with dynamic axon growth. bioRxiv, 2024–02.

66. Lebel, C., Walker, L., Leemans, A., Phillips, L., & Beaulieu, C. (2008). Microstructural maturation of the human brain from childhood to adulthood. Neuroimage, 40(3), 1044–1055.

67. McGee, A. W., Yang, Y., Fischer, Q. S., Daw, N. W., & Strittmatter, S. M. (2005). Experience-driven plasticity of visual cortex limited by myelin and Nogo receptor. Science, 309(5744), 2222–2226.

68. Hübener, M., & Bonhoeffer, T. (2014). Neuronal plasticity: beyond the critical period. Cell, 159(4), 727–737.

69. Mi, H., Muruganujan, A., Casagrande, J. T., & Thomas, P. D. (2013). Large-scale gene function analysis with the PANTHER classification system. Nature protocols, 8(8), 1551–1566.

70. Zhu, K., Bendl, J., Rahman, S., Vicari, J. M., Coleman, C., Clarence, T., … & Roussos, P. (2023). Multi-omic profiling of the developing human cerebral cortex at the single-cell level. Science Advances, 9(41), eadg3754.

