## Supplementary Figures for "Molecular mechanisms driving divergent development of the human frontal and visual cortex during prenatal development"

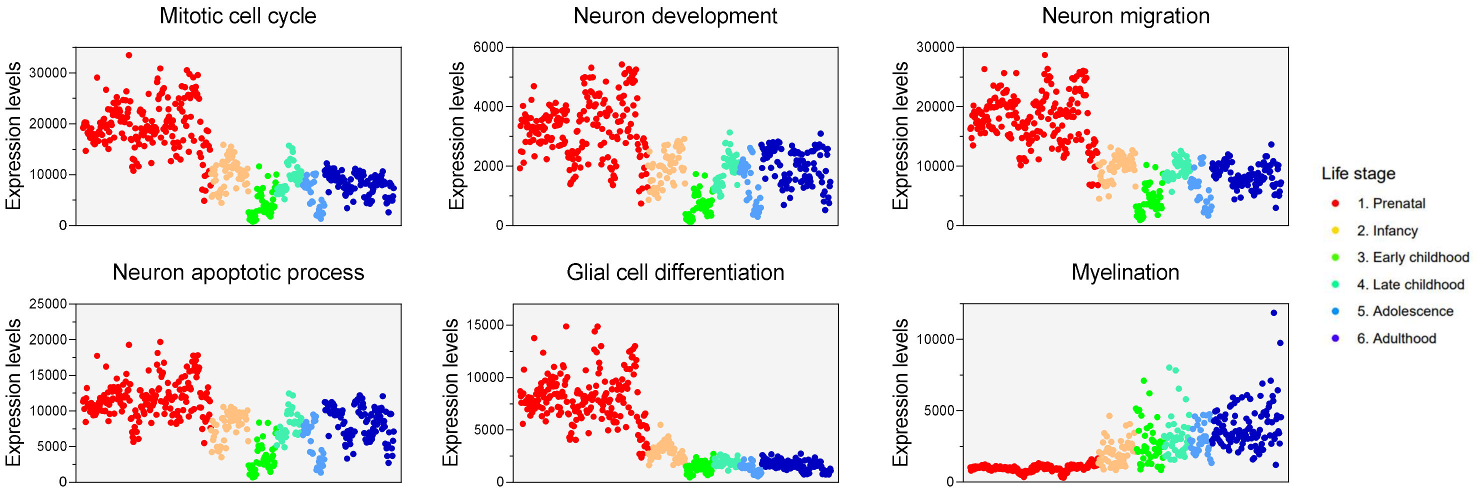


**Supplementary Figure 1.** Cumulative gene expression for selected pathways throughout the lifespan from 12 post-gestational weeks to 40 years. Gene expression of GO pathways follows the established developmental timeline of the processes. Each dot on the map is a brain region.

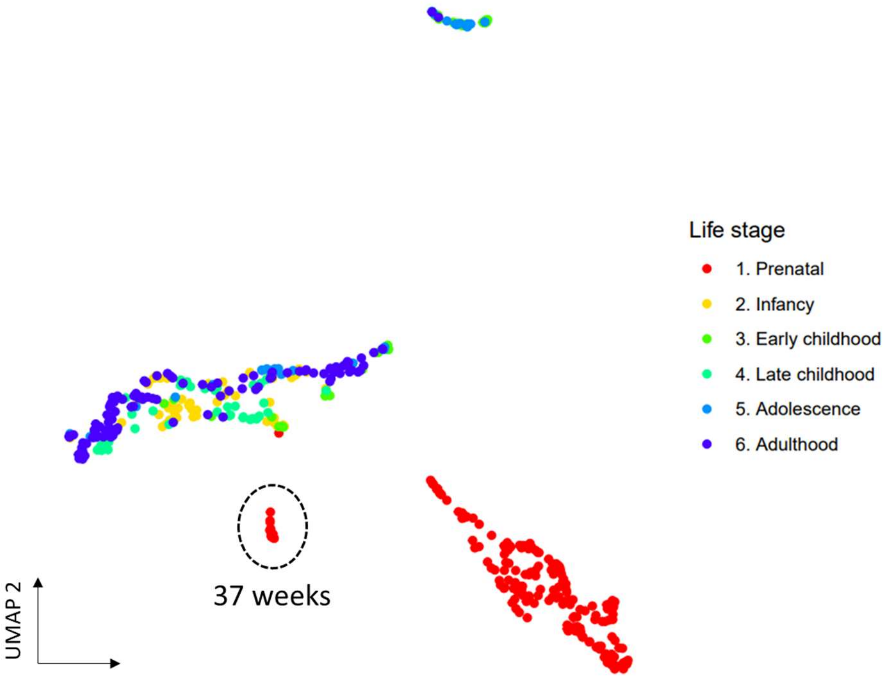


**Supplementary Figure 2.** A UMAP low-dimensional representation of gene expression coloured by life stages. Each dot represents a brain region. Brain regions at 37 post-gestational weeks demonstrate a marked transition in gene expression (represented between earlier prenatal samples and post-natal samples).
